# Tracking the fates of iron-labeled tumor cells in vivo using Magnetic Particle Imaging

**DOI:** 10.1101/2021.10.06.463387

**Authors:** Ashley V. Makela, Melissa A. Schott, Olivia C. Sehl, Julia J. Gevaert, Paula J. Foster, Christopher H. Contag

**Affiliations:** Institute for Quantitative Health Science and Engineering, Michigan State University, East Lansing MI USA; Western Michigan University, Kalamazoo MI USA; Department of Medical Biophysics, Robarts Research Institute, Western University, London ON Canada; Department of Biomedical Engineering, Michigan State University, East Lansing MI USA

**Keywords:** magnetic particle imaging, bioluminescent imaging, micron-sized iron oxide particles, iron-labeled cells, cell tracking, breast cancer, metastasis

## Abstract

The use of imaging to detect and monitor the movement and accumulation of cells in living subjects can provide significant insights that can improve our understanding of metastasis and guide therapeutic development. For cell tracking using Magnetic Resonance Imaging (MRI), cells are labeled with iron oxides and the effects of the iron on water provides contrast. However, due to low specificity and difficulties in quantification with MRI, other modalities and approaches need to be developed. Magnetic Particle Imaging (MPI) is an emerging imaging technique which directly detects magnetic iron, allowing for a specific, quantitative and sensitive readout. Here, we use MPI to image iron-labeled tumor cells longitudinally, from implantation and growth at a primary site to movement to distant anatomic sites. *In vivo* bioluminescent imaging (BLI) was used to localize tumor metastases and computed tomography (CT) allowed for correlation of these signals to anatomic locations. These three imaging modalities provide information on immune escape and metastasis of iron-labeled, and unlabeled, tumor cells, and the accumulation of cell-free iron contrast over time. We identified iron signals by MPI and tumor cells via BLI, and correlated these positive contrast images with CT scans to reveal the anatomic sites with cancer cells; histologic analysis confirmed the presence of iron-labeled tumor cells in the tissues, suggesting that the metastatic cells retained enough iron for MPI detection. The use of multi-modality cell tracking reveals the movement, accumulation and fates of labeled cells that will be helpful understanding cancer progression and guiding the development of targeted therapies.

---

*In vivo* cell tracking refers to imaging techniques which can be utilized to detect and monitor cells within the body. A widely-used imaging modality for tracking cells in living subjects is Magnetic Resonance Imaging (MRI). To detect cells by MRI, cells are typically labeled *ex vivo* using superparamagnetic iron oxide particles (SPIO) and are transferred into living subjects for tracking^1–3^. MRI detects SPIO indirectly, that is, the iron causes a distortion in the magnetic field, resulting in a region of signal void (black) within the MR image. The region of signal void encompasses an area much larger than the size of a cell, leading to very high sensitivity for counting labeled cells at a given anatomic site; and in fact, several studies have demonstrated that single cancer cells can be tracked using iron-based MRI^1–3^. A major limitation of this approach however, is the inability to directly quantify the amount of iron, and to quantify the number of cells at a given tissue site. Even though the number of discrete signal voids can be counted, area of signal void calculated, and the degree of contrast measured, it is not possible to determine cell numbers^4^.

Magnetic Particle Imaging (MPI) is an emerging imaging tool which directly detects SPIO. The MPI technique has been thoroughly described^5–7^, and is gaining acceptance as a preclinical imaging modality. In contrast to MRI images of iron, MPI detects superparamagnetic iron as a positive signal with little to no background, and appears as a ‘hot spot’, in other words yields positive contrast. Moreover, there is no signal attenuation from tissues, and therefore the MPI signal is linearly related to the amount of iron, allowing for direct quantification of iron and, with knowledge of the amount of iron per cell, the cell number can be estimated from images. This allows for specific, sensitive and quantitative imaging of cells, anywhere within the body.

For cancer cells the amount of iron per labeled cell decreases with cell division. In MRI this means that proliferating cells eventually contain too little iron to have an effect on the surrounding water protons, and these cells fall below the detection limit^1^. Where signals drop off for iron-labeled cells in MPI is not as well understood. If the proliferating cells remain within the MPI field of view the iron should still be detectable, however, significant dispersion of iron from the site reduces the cell density per voxel, leading some cells to fall below the intravoxel detection limit.

Another factor which may affect the detectability of cells by MPI is background signals associated with iron content in mouse chow. MPI signal due to mouse feed has been visualized in the region corresponding to the gastrointestinal (GI) tract in mice. When the region of interest is near the GI signal (ie. lower mammary fat pad (MFP) tumor), it can be difficult to distinguish the signal associated with iron-labeled cells from background. Tailored thresholding can overcome some of these issues, avoiding most of the GI signal. Further, the mice can be fasted prior to imaging, or the feed and bedding conditions can be changed to reduce signal.

Neither MRI or MPI can differentiate between live and dead cells. Additionally, MPI does not provide anatomic information. These limitations can be circumvented by the use of other imaging modalities. Since luciferases are enzymes requiring ATP and oxygen, they are dependent on cell metabolism, and as such, *in vivo* bioluminescent imaging (BLI) can be an indicator of cell viability. Luciferases can be engineered into cells of interest as an imaging reporter; the bioluminescent signals can be detected from an external vantage point and used as an indicator of cell viability. Computed tomography (CT) is a fast imaging technique which can provide the anatomic information required to understand where signal is located, within the primary tumor, the metastasis, and iron clearance from the body.

The purpose of this study was to evaluate patterns of tumor cell migration throughout the body, and use multimodality imaging employing MPI, BLI and CT and the resulting co-registered images to provide the information necessary to understand the location and accumulation of 1) live, iron-labeled cells (in the primary tumor or after metastasis), 2) live, non-labeled cells (proliferation) and 3) free iron label (cell death and uptake by other cells in the body).

## RESULTS AND DISCUSSION

The ability to track and quantify cells overtime in the bodies of living subjects can help us understand the fate and function of different cell types under a range of conditions. MPI can be used to track iron-labeled cells with high sensitivity, meaning that small numbers of iron-labeled cells can be detected. MPI directly detects superparamagnetic iron enabling visualization of the iron-labeled cells, and/or free iron-label, longitudinally. Pairing this new imaging technology with BLI provides an opportunity to monitor cell proliferation and cell movement to distant regions. CT was used in conjunction with these imaging methods to provide anatomic references for the signal. Each of this imaging modalities provided 3-dimensional data sets to better understand exactly where the signals localized within the body. MPI allowed for tracking of iron-labeled cells, initially as they formed a primary mammary carcinoma, as the cells metastasized and later associated with iron-label clearance.

### MPIO cell labeling and MPI characterization

Cell pellets (n = 3) containing 7.5×10^5^ cells were imaged by MPI and quantified to determine an average total iron content of 12.05 +/- 1.42 μg Fe, corresponding to an average of 16.07 pg Fe/cell. 100% of the cells were seen to be labeled by micron-sized iron oxide particles (MPIO), to varying degrees (**Figure 1a**). The MPIO-labeled 4T1BGL cells can be visualized under fluorescence as green (GFP), with blue MPIO within the cytoplasm (**Figure 1b**). Relaxometry was then used to investigate the performance of the MPIO as a free iron agent (1 μg), suspended in PBS or as intracellular MPIO (3×10^5^ cells, n = 3; **Figure 1c**). The amplitude of the signal per ng Fe was the same for intracellular vs free MPIO (11.65 +/- 5.14 vs 7.82 +/- 0.59 signal (a.u.)/ng Fe, *p =* 0.2689). The full width at half max (FWHM), corresponding to spatial resolution, of intracellular vs free MPIO were also not significantly different (39.22 +/- 3.39 vs 31 +/- 7.2 mT respectively, *p =* 0.1485). These values correspond to a spatial resolution of 6.88 mm and 5.44 mm, respectively. The variance in the performance of the free MPIO samples appears to be larger than that of the intracellular MPIO.

**Figure 1.**
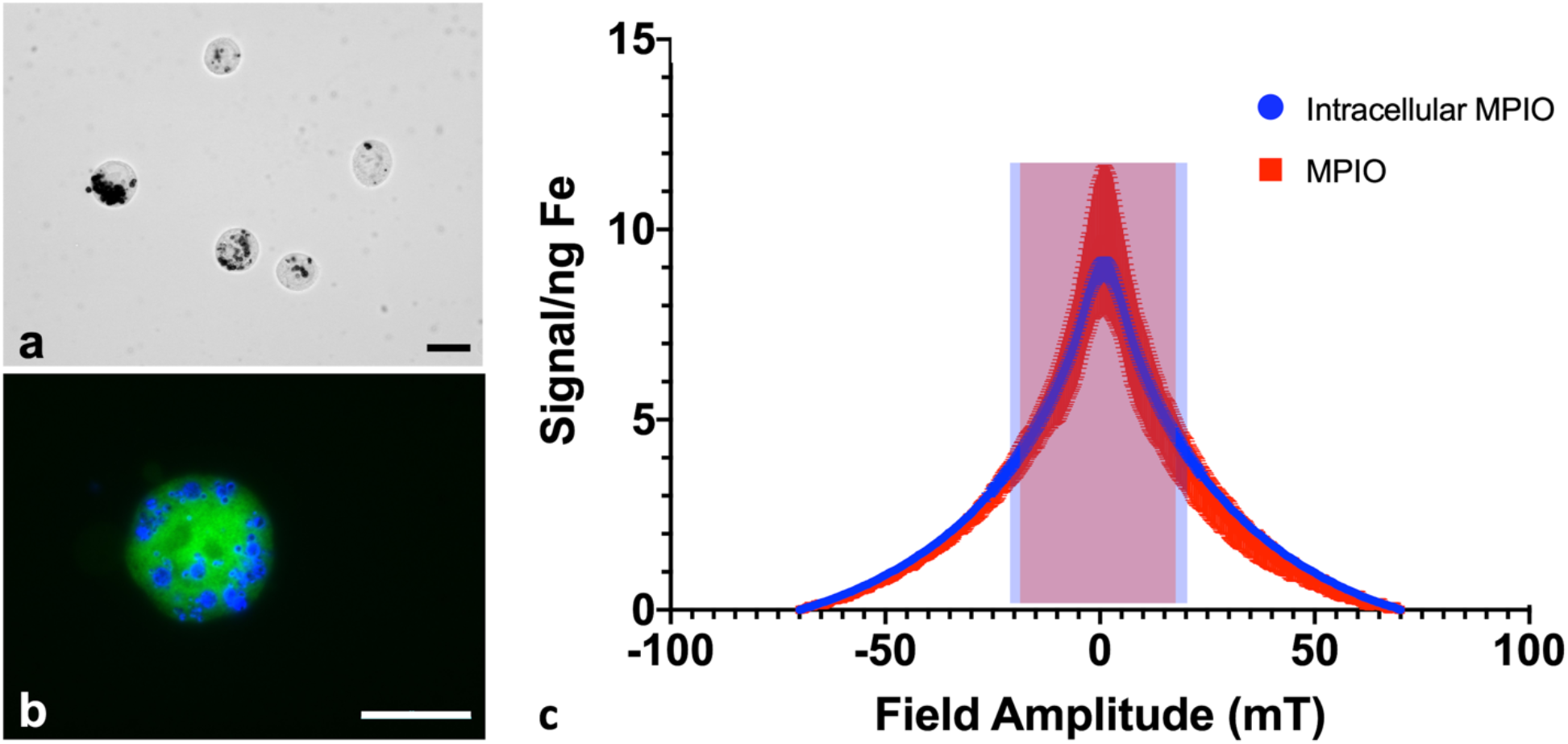
Labeling and MPI performance of free MPIO vs MPIO-labeled cells. (a) Brightfield microscopy can identify MPIO (black) within the 4T1BGL cells. (b) Fluorescent microscopy visualizes MPIO particles (blue) within 4T1BGL cells (green). (c) Relaxometry displays the signal amplitude (signal/ng Fe) and resolution (FWHM) of intracellular (shaded blue) vs free (shaded red) MPIO. Each data point is displayed as +/- std dev. Scale bar = 20 μm. MPI: magnetic particle imaging; MPIO: micron-sized iron oxide particles; FWHM: full width at half max.

The 4T1 cancer cells could be easily labeled through coincubation with MPIO. Although it’s unlikely there was detection of single cells with MPI, as seen using MRI^2,3^, it is possible that MPI could detect cells even with reduced iron content per cell, resulting from cell proliferation. With MRI, the ability to detect daughter cells with a reduced amount of iron within them, is diminished. Because MPI relies on the detection of iron itself, MPI could still quantify the amount of iron in a region of interest, even after some loss of iron label in an individual cell. However, this change in intracellular iron convolutes any quantification with respect to cell number; the original amount of iron loading could not be used to determine cell number.

The signal strength of the MPIO was not altered by cellular internalization, which is in agreement with Melo et al^8^. Others have reported a loss in signal with internalization for other SPIO^9^. These different findings may be due to the differences in composition between the MPIO and SPIO particles; the MPIO is embedded in a polymer which may prohibit a change in the relaxation properties when exposed to different environments when compared to the SPIO. Differences in composition may also have an effect on MPI resolution. While we identified no significant change in resolution between 1.63 μm, encapsulated MPIO and MPIO-labeled cells, Melo et al^8^ found there was a decrease in resolution when 0.9 μm, unencapsulated MPIO were internalized into cells. Although we did detect some differences in resolution we do not expect this finding to lead to inaccurate quantification whether the MPIO is still within the cell or being cleared from the body.

### Multimodal *in vivo* imaging

3D MPI was used to image and quantify iron content at every time point within the primary tumor (**Figure 2a**) and throughout the mouse. The amount of iron in the tumor at day (d)0 iron was calculated to be 4.69 +/- 0.45 μg, resulting in an estimate of 15.64 pg Fe/cell, which was similar to the *in vitro* quantification of cell pellets (*p* = 0.7175). The amount of iron measured in the primary tumor decreased over time presumably due to cell death, and subsequent release of the iron-label, or to a lesser extent, escape of iron-labeled cells^10^. Iron amounts within the primary tumor were: d0 – 4.69 +/- 0.45 μg, d5 – 3.55 +/- 0.41 μg, d10 – 3.15 +/- 0.24 μg, d17 – 2.49 +/- 0.33 μg and d26 – 1.994 +/- 0.4 μg. Iron content was normalized to d0 for each tumor. The relative iron amounts were: d0 – 100%, d5 – 76.5%, d10 – 52.34%, d17 – 51.33% and d26 – 41.47% (**Figure 2b**). These values correspond to an average loss of 1.6% of the original iron amount per day.

**Figure 2.**
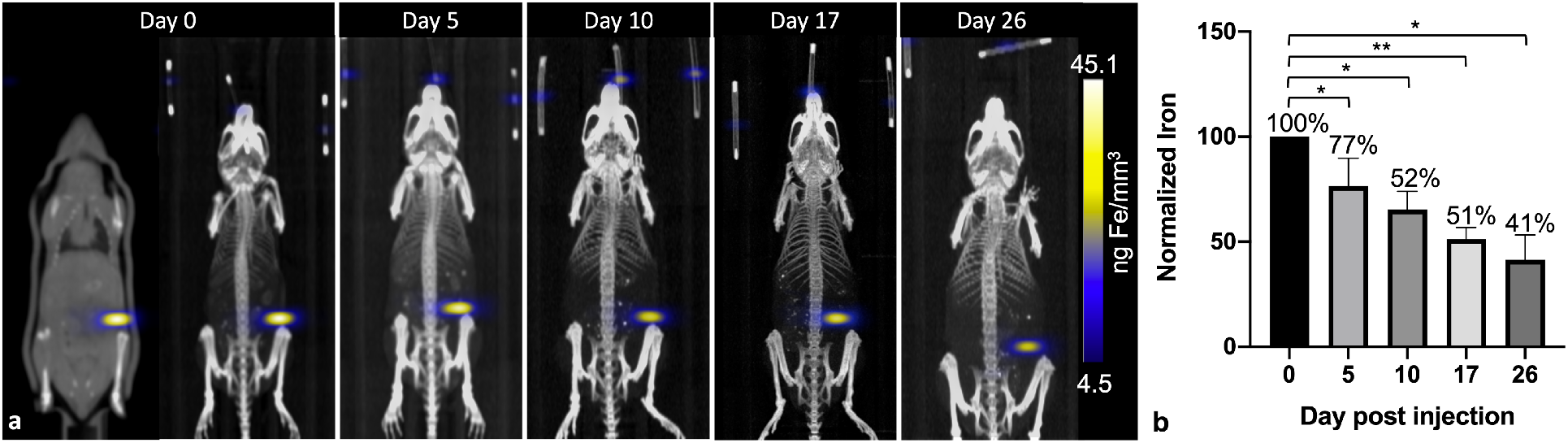
*In vivo* longitudinal MPI to visualize the primary tumor. Visually, the signal intensity in each primary tumor decreases from d0 to d26 (a). The amount of iron was quantified using total MPI signal and normalized to d0; the iron quantified in the primary tumor decreases up to d26 (b). \**p*<0.05, \*\**p*<0.01. MPI = magnetic particle imaging

MPI signal was also identified in regions outside of the primary tumor throughout the mouse, which correlated with the liver and lymph nodes, as identified by CT (**Figure 3**). When the iron in these regions were quantified, the total amount of iron accounted for, including the primary tumor, was then d5 = 93.04%, d10 = 79.65%, d17 = 67.91% and d26 = 63.65%. There were regions of MPI signal which were below the quantification threshold and these were not including in these calculations.

**Figure 3.**
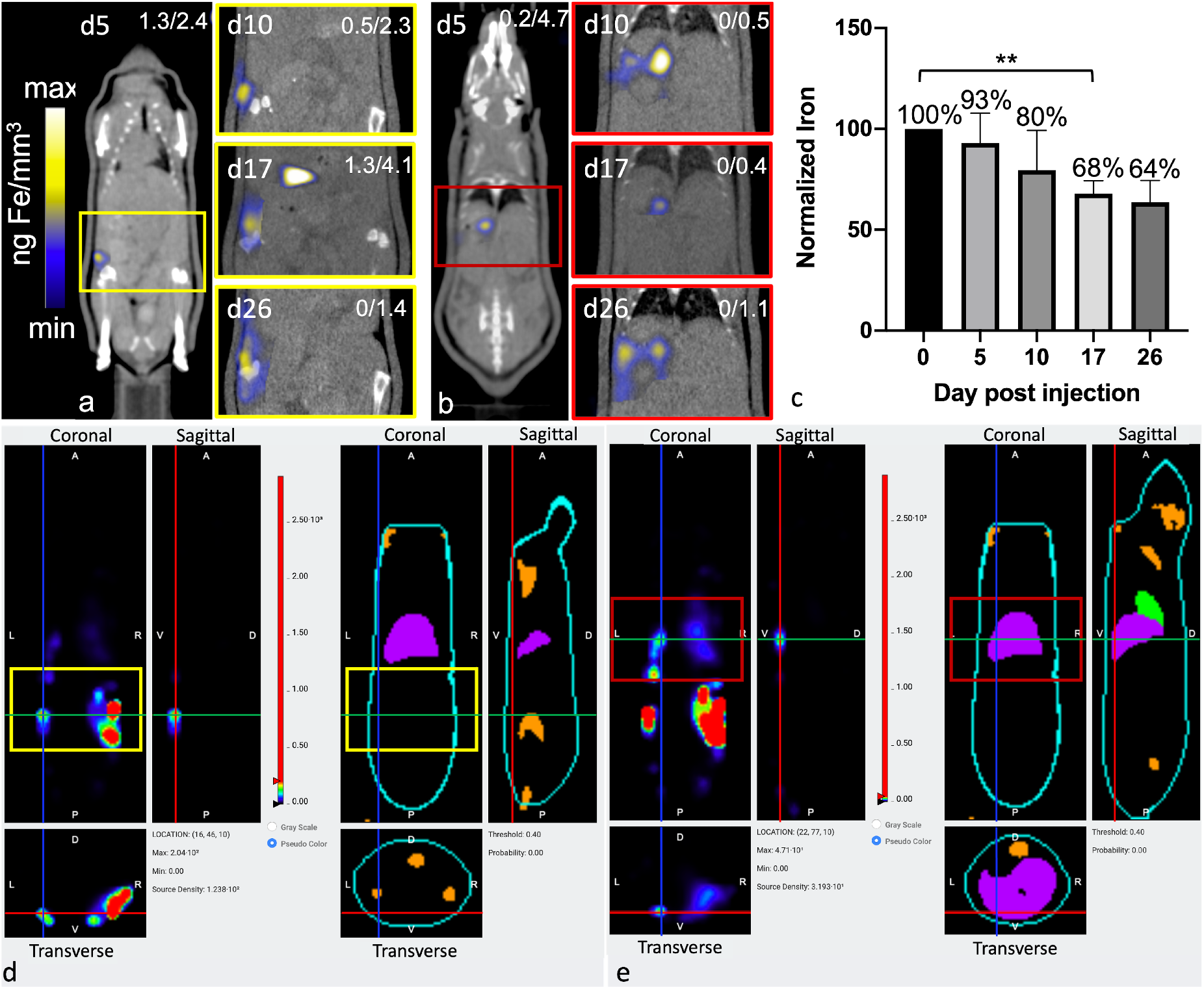
Extratumoral iron and metastases detected longitudinally by MPI and BLt. Regions of iron accumulation were noted in the same mouse in the liver (a) or lymph nodes (b), beginning at d5 to d26. The amount of iron in all extratumoral regions were quantified using total MPI signal. The total whole body iron content (primary tumor + extratumoral) for all time points was normalized to d0 (c). BLt identifies luminescence from the same region as identified in MPI, at d26 (lymph node, d - yellow and liver, e - red). MPI CLUT identifies numerical max/min values of ng Fe/mm^3^ for each image (top, right). **p<0.01. MPI, magnetic particle imaging; BLt, bioluminescent tomography; CLUT, color lookup table

The free iron-label could be tracked to the liver and to lymph nodes; these tissues have many phagocytic cells which are used for elimination of foreign debris^11^. However, the composition of MPIO particles does not allow for biodegradation and therefore these particles cannot be cleared from the body^12^. When the total iron content (primary tumor + distant regions) was calculated, up to 80% of the initial iron content could be accounted for, up to d10. At d26 only 64% of the original iron content could be identified, which could be due to the spread of small amounts of the iron throughout the subject which was below the intravoxel detection threshold. It is important to note that MPIO particles are not optimized for MPI and therefore, detection of cells labeled with a more MPI tailored SPIO could provide the means to track smaller numbers of cells to these extratumoral regions.

BLI was used to visualize bioluminescence from the 4T1BGL cells which were initially injected into the MFP on d0, up until d26 in both the primary tumor site and metastases. BLI signal was present in the primary tumor site from d0 to d26. BLI signal was also noted in extratumoral regions, including the lymph nodes, lungs and liver, presumably as metastases, beginning at d5. Some of these regions did not have any MPI-detectable iron associated with them, indicating that there is no iron present or it is below the MPI detection threshold.

CT images were used to correlate the BLI or MPI signals to an anatomic region. BLI and MPI signal were identified in the same region in some cases, suggesting the presence of live MPIO-labeled 4T1BGL cells and/or unlabeled 4T1BGL cells with other MPIO-labeled cells in the immediate surrounding region, for example, macrophages.

### Fasting to reduce gastrointestinal MPI signal

Decreasing the MPI signal in the GI region would improve imaging of nearby regions of interest. Combinations of treatments, including mouse fasting, changes in bedding (cotton or corn), and use of laxatives, were applied in attempt to reduce MPI signal in the GI region. All mice showed a reduction in MPI signal from GI region after fasting (**Figure 4**). When mice were housed with cotton bedding and fasted, MPI signal was reduced by 36% whereas with corn bedding, further reduction in MPI signal was observed (86%). With laxatives and fasting, GI signal was reduced by 35% with cotton bedding and 79% with corn bedding. Therefore the addition of laxatives had less of an impact on GI signal than the type of cage bedding.

**Figure 4.**
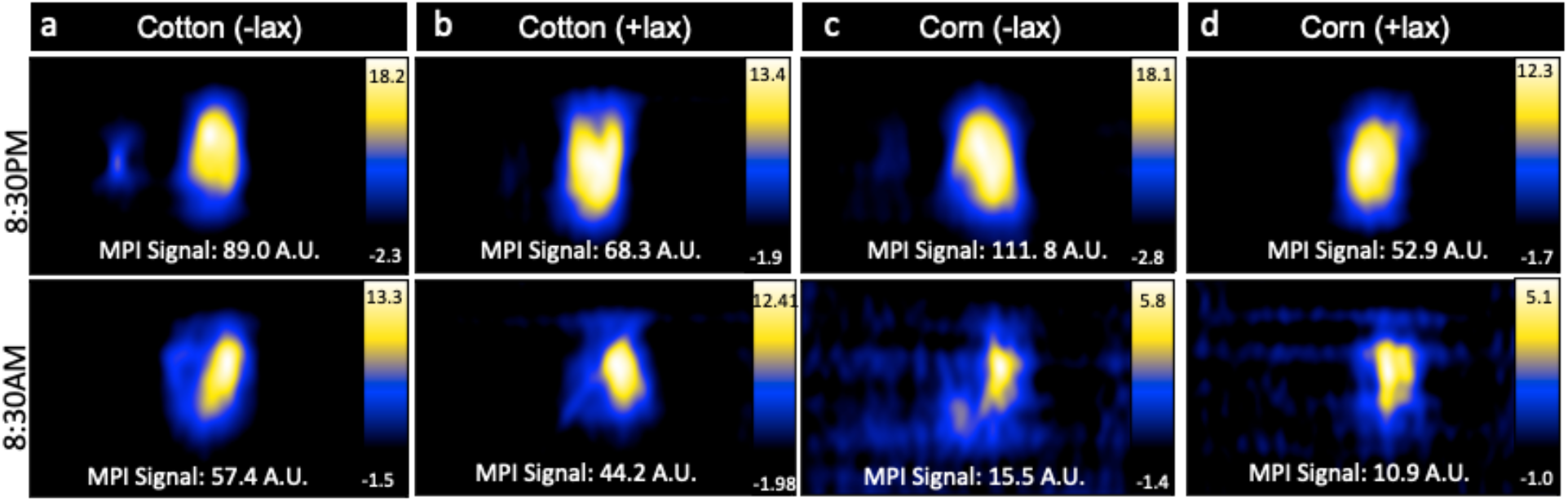
MPI of mice fasted for 12h overnight using either cotton or corn bedding, with and without laxatives. Gastrointenstinal (GI) MPI signal was reduced after fasting in all mice. Mice in cages with corn bedding had a lower GI signal compared to cotton bedding post-fast. Laxatives did not appear to impact GI signal. MPI, magnetic particle imaging.

### Histology

Within tumor and lymph node tissue, both iron (blue)-labeled and unlabeled 4T1BGL cells (green) were identified (**Figure 5**). These were seen within the central region of the tumor, and within both micrometastases and as single cells in the lymph nodes. F4/80+ cells (macrophages: red) were also identified in both tissues (**Figure 6**). Perls’ Prussian Blue (PPB) staining on adjacent histology sections identifies the iron as blue within the tissue. Within tumors, F4/80+ cells were seen around the periphery with no PPB staining (MPIO) present; macrophages are commonly found in this invasive region^13,14^. Within the lymph node, both MPIO-labeled and unlabeled F4/80+ macrophages were present.

**Figure 5.**
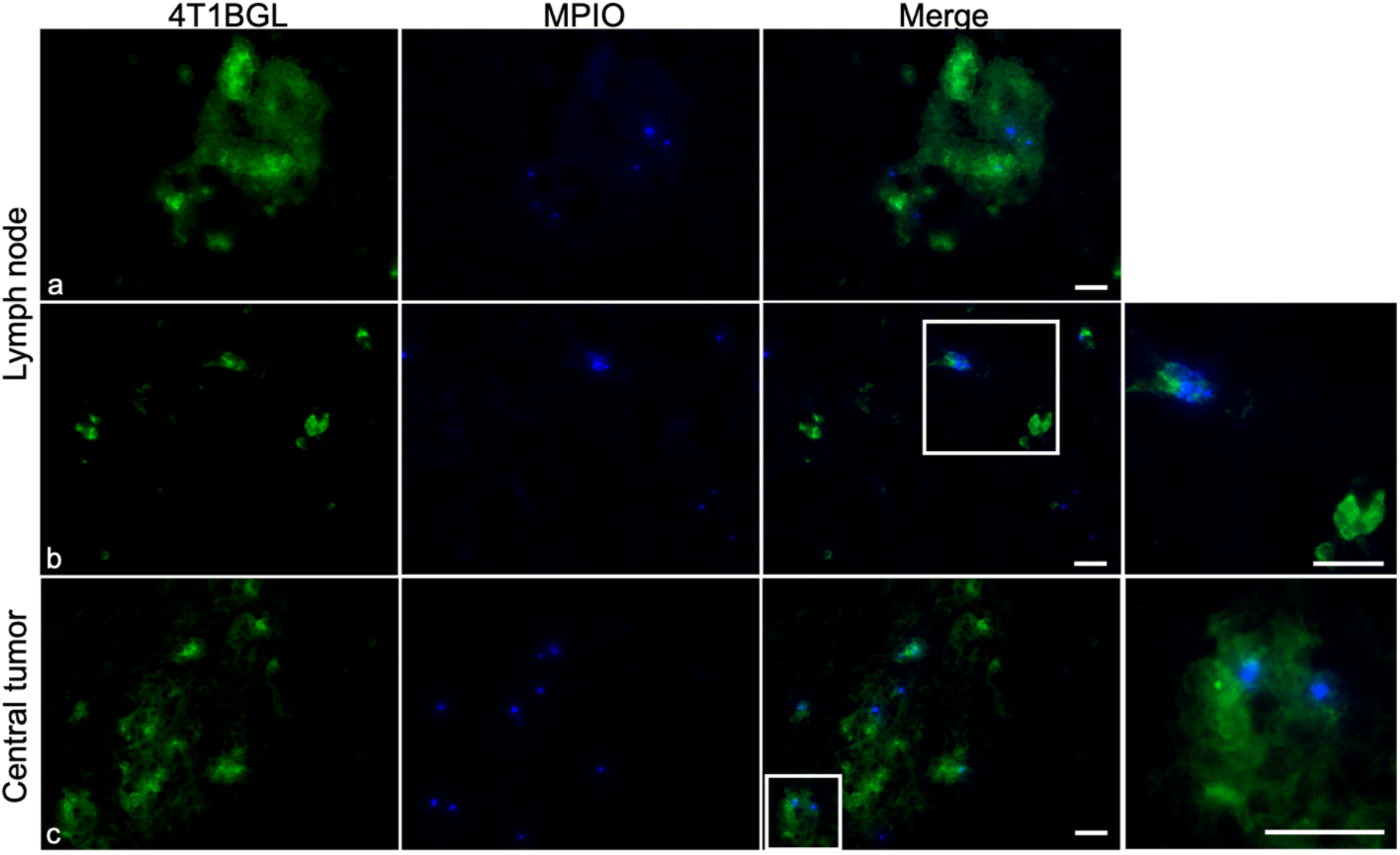
Histological analysis of excised lymph nodes and 4T1BGL tumors. Fluorescence microscopy detected 4T1BGL cells (green) and MPIO (blue) in micrometastases (a) and single cells (b) in the lymph nodes and within the central region of the tumor (c). Fluorescence from the 4T1BGL cells and MPIO were located within similar regions, relating to the presence of MPIO-labeled 4T1BGL. There was the presence of 4T1BGL cells which did not have any corresponding MPIO signal, suggesting these cells had already lost the MPIO label. Zoomed insets (far right) highlight the heterogeneity of MPIO-labeled and unlabeled 4T1BGL cells in each location. Scale bar = 20 μm, MPIO = micron-sized iron oxide particles.

**Figure 6.**
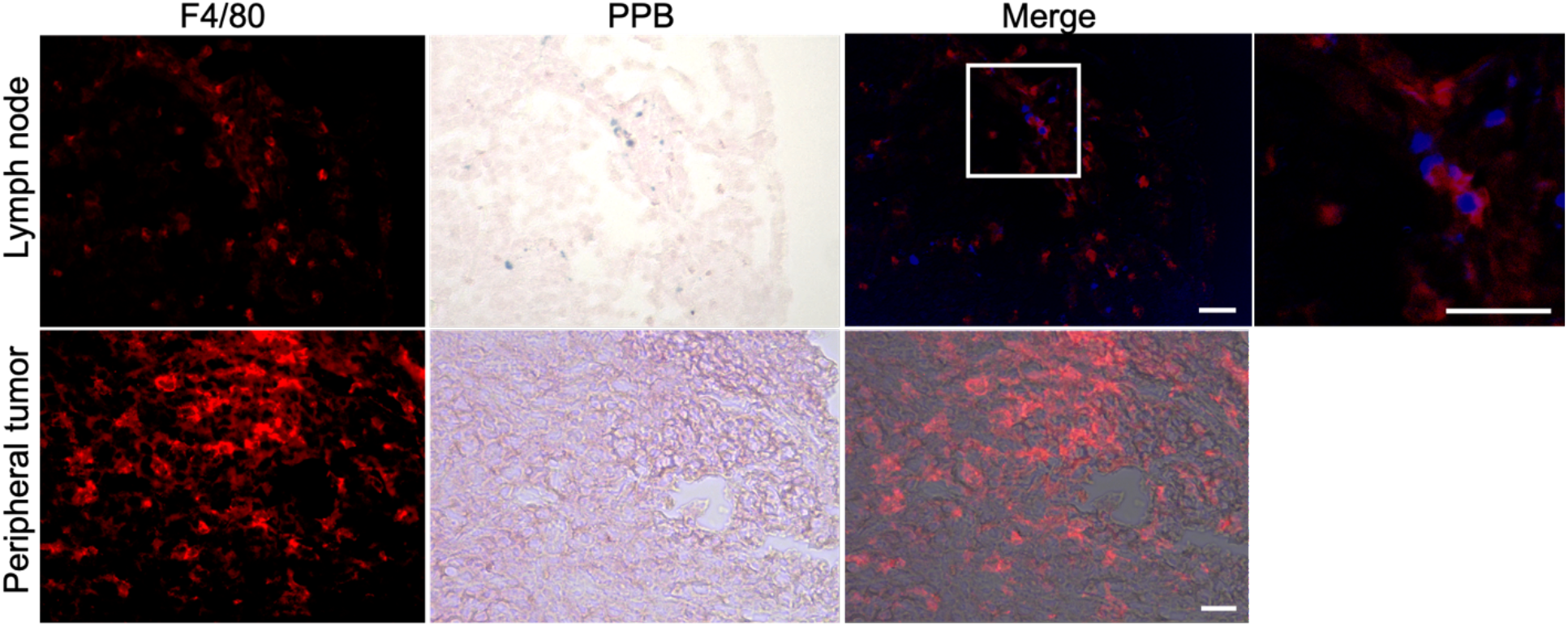
Histological analysis of excised lymph node and 4T1BGL tumor. Macrophages (F4/80+; red) were identified in the lymph nodes, corresponding to the location of the MPIO (Perls Prussian Blue, PPB). PPB+ cells were colorized blue for merge with fluorescent F4/80. Macrophages were present in the periphery of the tumor and did not correspond to the presence of MPIO. Scale bars = 20 μm. MPIO = micron-sized iron oxide particles.

Macrophages are natural scavengers which take up foreign debris, such as the free MPIO. Macrophages can be present in metastases, known as metastasis-associated macrophages^15^ but phagocytic cells are also present in the liver (ie Kupffer cells) and lymph nodes. Histological examination of these regions confirmed that the original, MPIO-labeled 4T1BGL cells were present in the central region of the tumor, with unlabeled macrophages populating the outer periphery. Lymph nodes contained unlabeled 4T1BGL cells, MPIO-labeled 4T1BGL cells and macrophages which had taken up the MPIO. These results suggest that the iron remaining in those cells that metastasized could contribute to MPI signal as they travel to remote regions.

## Conclusions

This study presented the use of MPI, BLI and CT as a multi-modality imaging approach to understand the metastatic process, originating from a primary tumor. MPI was utilized to track iron present as MPIO-labeled 4T1BGL, MPIO-labeled phagocytic cells and free MPIO-label. BLI identified regions in which there were viable 4T1BGL tumor cells and CT provided anatomic reference for each of these imaging modalities. Together, these three imaging modalities allowed for a better understand of the presence of metastases and clearance of the free MPIO-label, due to tumor cell death.

## METHODS

### Cell Culture

Murine mammary carcinoma cells of BALB/c origin (4T1) were built to stably express Blasticidin resistance, *Luciola Italica* luciferase (Red-FLuc) and green fluorescent protein (GFP) (4T1BGL) using the *Sleeping Beauty* transposon^16^. Cells were maintained in RPMI containing 10% FBS and penicillin/streptomycin at 37°C and 5% CO_2_, with Blasticidin selection. For iron-labeling, 3×10^6^ 4T1BGL cells were plated and 24 hours (h) later incubated with 25 μg/mL of MPIO (1.63 μm in diameter, Carboxy Encapsulated Magnetic Fluorescent Polymer, conjugated with Glacial Blue, Bangs Laboratories Inc, IN, USA) overnight. Cells were then washed three times with PBS, collected after trypsinization, and washed three more times with PBS to remove residual unincorporated MPIO prior to *in vitro* evaluation or injection into animals.

### *In vitro* studies

MPIO-labeled 4T1BGL cells were pelleted as triplicates of 7.5×10^5^ cells for imaging by MPI. MPI was acquired using a Momentum MPI system (Magnetic Insight Inc, Alameda, USA; MSU location unless otherwise stated) using a 2D projection scan with default setting (5.7 T/m gradient, 16.5 mT (X-channel) and 17 mT (Z-channel) excitation). The imaging parameters were: FOV = 4 × 6 × 6 cm, 1 average and acquisition time of 15 seconds (s). Iron quantified from this acquisition was used to estimate the amount of iron per cell.

Particle relaxometry was performed on MPIO-labeled 4T1BGL pellets (3×10^5^, n = 3) and free MPIO (1 μg) to investigate any changes in MPIO signal due to cellular internalization. Signal amplitude (peak signal strength) and spatial resolution (calculated using FWHM) were calculated by averaging the values obtained by the positive and negative scans.

### Animal Model

Female BALB/c mice (6-8 weeks; Charles River USA) were obtained and cared for in accordance with the standards of Michigan State University Institutional Animal Care and Use Committee. Mice were anesthetized with isoflurane administered at 2% in oxygen followed by an injection of 300,000 MPIO-labeled 4T1BGL cells (n = 6) (>90% viability, measured using the trypan blue exclusion assay) suspended in 50 μl PBS into the 4^th^ (inguinal) MFP, as previously reported^17^.

### *In vivo* iron-labeled 4T1BGL imaging protocol

BLI, MPI and CT imaging studies were performed on mice at day (d) 0 (n = 6), 5 (n = 6), 10 (n = 5), 17 (n = 4) and 26 (n = 3) post implantation (pi) of the 4T1BGL cells. Mice were imaged in a body conforming animal mold (BCAM; InVivo Analytics Inc, NY, USA) which could be transported to each imaging modality without moving the mouse. Iron fiducial markers were placed in 3 dimensions to allow for co-registration.

Mice received an intraperitoneal injection of D-luciferin (150 mg/kg, PerkinElmer, MA, USA) 10-15 minutes prior to beginning the imaging protocol. First, bioluminescent tomography (BLt) was performed using an IVIS Spectrum system (PerkinElmer). Mice within the BCAM were placed into a mirror gantry (InVivo Analytics Inc) situated in the IVIS. Images were captured as previously described^18^ to enable the reconstruction of the imaging data to a 3D data set.

The BCAM was then transferred to a MPI bed adaptor (InVivo Analytics Inc) without moving the mouse. 3D MPI tomographic scans were performed using the following parameters: FOV: 12 × 6 × 6 cm, 5.7 T/m gradient, 16.5 mT (X-channel) and 17 mT (Z-channel) excitation, 35 projections and 1 average with an acquisition time of ∼30 mins.

The BCAM was then transferred to a Quantum GX microCT scanner (PerkinElmer). Whole body CT images were acquired using 3 × 8s scans with the following parameters: 90 kV voltage, 88 μA amperage, 72 mm acquisition FOV and 60 mm reconstruction FOV resulting in 240 μm voxels.

### Image analysis of cell pellets and *in vivo* studies

MPI data sets were visualized and analyzed utilizing Horos imaging software (Horos is a free and open source code software program that is distributed free of charge under the LGPL license at Horosproject.org and sponsored by Nimble Co LLC d/b/a Purview in Annapolis, MD USA). To quantify MPI signal from 3D datasets in the primary tumor, cell pellets, and fiducials, a threshold was set to capture signal above 10x standard deviation (sdev) of the noise. The high threshold value was necessary to avoid inclusion of GI signal present in the animal, due to close proximity to the primary MFP tumor. When quantifying extratumoral iron, pixels under 2x sdev of the noise were set to 0, allowing for visualization and precise capture of the signals above this threshold. The low threshold value allowed for identification of small amounts of signal which were commonly in regions distant from GI signal. Sdev of the noise was determined using a slice from the 3D dataset which did not include any of the primary tumor or fiducial markers.

Regions of signal from 3D images were automatically thresholded, slice by slice, creating a 3D volume. Total MPI signal was calculated by *mean signal × volume*. The signal intensities using the color look up table were converted to iron concentration (Fe/mm^3^), based on signal from the iron fiducials.

Calibration curves were created by imaging different amounts of iron and then plotting the known amount of iron to the total MPI signal generated, using matched threshold values and imaging parameters (such as 2D or 3D image acquisition and scan mode), dependent on acquisition parameters of data set being analyzed. Iron content in cell pellets and *in vivo* was determined by substituting the total MPI signal from the ROI into the calibration curve equation with the intercept set to 0.

3D BLt dicoms were visualized using InVivoPLOT (InVivo Analytics Inc) and exported for co-registration in Horos imaging software. The signal from the BLt images was compared to that of MPI and anatomical location was determined using CT.

### Fasting protocol and MPI acquisition

NOD SCID gamma mice were bred at Robarts Research Institute and cared for in accordance with the standards of Canadian Council on Animal Care, under an approved protocol by the Animal Use Subcommittee of Western University’s Council on Animal Care. Mice were fasted overnight for 12h in cages with only water and bedding (either cotton or corn). All other cage elements were removed. Four fasting conditions were tested (n = 1 per condition): (i) cotton bedding without laxative (ii) cotton bedding with laxative (iii) corn bedding without laxative and (iv) corn bedding with laxative. One tablet of chocolate laxative was left in the cage for conditions (ii) and (iv). After fasting and imaging, mice were returned to fully restored cages.

MPI fasting data was acquired at Robarts Research Institute, Western University using 2D high sensitivity setting (3.0 T/m gradient, 20 mT (X-channel) and 26 mT (Z-channel) excitation. The imaging parameters were: FOV = 12 × 6 × 6 cm, 1 average and acquisition time of 2.8m. Mice were scanned individually at two time points; immediately prior to fasting, and again 12h later. Total MPI signal was quantified by setting a threshold to capture signal above 5x sdev of the noise.

### Histological Analysis

After *in vivo* imaging, tumors and lymph nodes were removed and fixed in 4% paraformaldehyde for 24h followed by cryopreservation through serial submersion in sucrose (10%, 20% and 30%). Sections were then frozen in optimal cutting temperature compound. Tissue was sectioned using a cryostat (10 μm sections). Sections were then stained with PPB for iron and primary F4/80 Monoclonal Antibody (BM8), (eBioscience™ from Thermo Fisher Scientific, catalog #14-4801-85) with Goat anti-Rat IgG (H+L) Cross-Adsorbed Secondary Antibody, Alexa Fluor 647 (Thermo Fisher Scientific, catalog #A-21247). The 4T1BGL cells could be identified as green (GFP) and MPIO particles as blue. A sample of fixed MPIO-labeled cells were visualized under brightfield and fluorescence microscopy to confirm labeling of cells.

Microscopy was performed using a Nikon Eclipse Ci microscope equipped with a Nikon DS-Fi3 high-definition camera (Nikon Instruments Inc. Tokyo, Japan) for fluorescent imaging and CoolSNAP DYNO (Photometrics, AZ, USA) for color and brightfield acquisition.

### Statistical Analysis

Statistical analyses were performed using Prism software (8.0.2, GraphPad Inc., CA, USA). Cell pellets vs d0 iron content and relaxometry data were analyzed using an unpaired t-test. Percent of iron at each time point was analyzed using a mixed-effects analysis with Geisser-Greenhouse correction, followed by Tukey’s multiple comparisons test. Data are expressed as mean +/- standard deviation; *p*<0.05 was considered a significant finding.

## AUTHOR INFORMATION

### Author Contributions

The manuscript was written through contributions of all authors. All authors have given approval to the final version of the manuscript.

### Notes

Dr. Christopher Contag is on the advisory board for InVivo Analytics Inc.

## ACKNOWLEDGMENT

We acknowledge and thank InVivo Analytics for the use of the tools and technology to perform BLt and use of the software InVivoPlot. The authors would like to that the James and Kathleen Cornelius Endowment for the support that enabled this study.

## ABBREVIATIONS

MRI: magnetic resonance imaging
MPI: magnetic particle imaging
BLI: bioluminescent imaging
CT: computed tomography
SPIO: superparamagnetic iron oxide particles
GI: gastrointestinal
MFP: mammary fat pad
MPIO: micron-sized iron oxide particles
FWHM: full width at half max
BLt: bioluminescence tomography
PPB: Perls’ Prussian Blue
GFP: green fluorescent protein
pi: post implantation
BCAM: body conforming animal mold
sdev: standard deviation

